# Combining high-density electromyography and ultrafast ultrasound to assess individual motor unit properties *in vivo*

**DOI:** 10.1101/2023.07.03.547503

**Authors:** M. Carbonaro, R. Rohlén, S. Seoni, K. M. Meiburger, T. Vieira, C. Grönlund, A. Botter

**Affiliations:** LISiN, Department of Electronics and Telecommunication, Politecnico di Torino, Turin, Italy; PoliToBIOMed Lab, Politecnico di Torino, Turin, Italy; Department of Biomedical Engineering, Lund University, Lund, Sweden; Department of Radiation Sciences, Radiation Physics, Biomedical Engineering, Umeå University, Umeå, Sweden; Biolab, Department of Electronics and Telecommunications, Politecnico di Torino, Turin, Italy

**Keywords:** high-density surface emg, ultrafast ultrasound, motor unit, averaging, independent component analysis

## Abstract

This study aims to compare two methods for the identification of anatomical and mechanical motor unit (MU) properties through the integration of high-density surface electromyography (HDsEMG) and ultrafast ultrasound (UUS). The two approaches rely on a combined analysis of the firing pattern of active MUs, identified from HDsEMG, and tissue velocity sequences of the muscle cross-section, obtained from UUS. The first method is the spike-triggered averaging (STA) of the tissue velocity sequence based on the occurrences of MU firings. The second is a method based on spatio-temporal independent component analysis (STICA) enhanced with the information of single MU firings. We compared the capability of these two approaches to identify the regions where single MU fibers are located within the muscle cross-section (MU displacement area) *in vivo*. HDsEMG signals and UUS images were detected simultaneously from biceps brachii in ten participants (6 males and 4 females) during low-level isometric elbow flexions. Experimental signals were processed by implementing both STA and STICA approaches. The medio-lateral distance between the estimated MU displacement areas and the centroid of the MU action potential distributions was used to compare the two methods. We found that STICA and STA are able to detect MU displacement areas. However, STICA provides more precise estimations to the detriment of higher computational complexity.

## I. INTRODUCTION

HIGH density surface electromyography (HDsEMG) and ultrafast ultrasound (UUS) provide complementary information about electrical and mechanical muscle properties. Both techniques have been proven successful in characterising single motor unit (MU) properties, opening exciting fronts in studying the neural control of muscle contraction from different perspectives. While HDsEMG decomposition enables identifying the firing pattern of individual MUs and assessing the characteristic MU action potentials (MUAP, electrophysiological information) [1], [2], the analysis of cross-sectional tissue velocity sequences extracted from UUS [3] can provide anatomical and mechanical characteristics of MU fibers [4]–[6].

Recently, these two techniques have been combined to exploit their complementarity and provide a comprehensive description of the neuromechanical MU properties [7]–[10]. This integration uses single MU firing instants, identified from HDsEMG decomposition, to isolate the corresponding muscle tissue mechanical response from UUS tissue velocity sequences (Fig. 1). In this way, it is possible to assess movements in the physiological cross-sectional area of the muscle associated with the excitation of single MU (hereafter referred to as *MU displacement area*) and their twitch characteristics. In this regard, two possible approaches can be implemented. The first one is the spike-triggered averaging (STA) of the tissue velocity sequence based on the occurrences of individual MU firings, as proposed for single MU analysis of mechanomyograms [11]. The second one, recently proposed by our group, is to decompose the tissue velocity sequences with spatio-temporal independent components analysis (STICA) [12] and use the firing information from HDsEMG to select, among the identified components, those associated with MU activity [7].

**Fig. 1:**
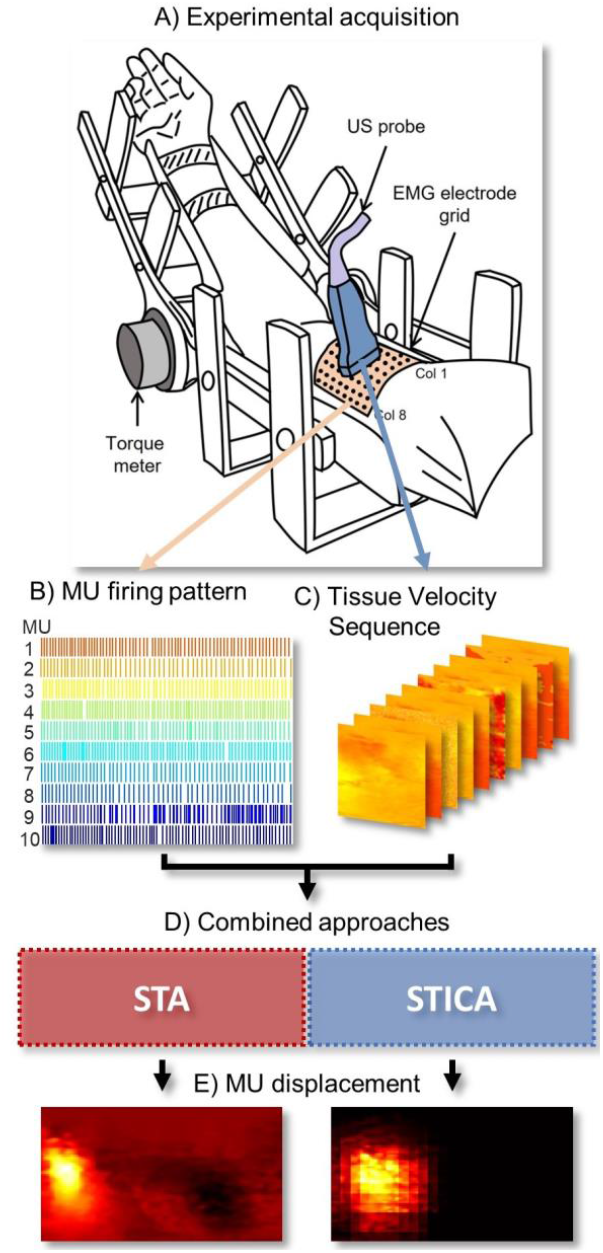
Overview of data detection and analysis. A) Simultaneous acquisition of HDsEMG and ultrafast ultrasound data during isometric contractions. B) Example of a firing pattern of active MUs decomposed from HDsEMG signals. C) Estimated image sequence of the displacement velocities of the muscle tissue. D) Combined analysis of the MU firing pattern and tissue velocity sequence with two approaches, one based on the spike triggered averaging (STA) and one based on spatio-temporal independent component analysis (STICA). E) Output of the algorithms: images of muscle tissue displacement areas related to a single MU activation.

Previously, we used an electromechanical computer model to test the capability of the two methods to correctly identify single MU displacement areas in simulated contractions with different degrees of neural excitation (i.e., the number of active MUs) and different levels of MU synchronisation (i.e., degree of dependency between firing instants of different MUs) [13]. These variables and the duration of both EMG and ultrasound time series are potentially critical factors impacting both methods. Indeed, we showed that the performance of both approaches was negatively affected by the number of active MUs and synchronisation levels. However, STICA provided a more robust estimation of the MU displacement areas under all the tested conditions.

This study aimed to test whether these *in-silico* results extent to experimental conditions during isometric elbow flexions. Since the location of the MU fibers within the muscle cross-section is unknown in experimental data, we quantified the agreement between the HDsEMG amplitude distribution of the MUAP and the location of the corresponding MU displacement area identified by applying STA and STICA to UUS image sequences.

## II. Methods

Ten participants (mean ± SD, 29.2 ± 4.6 years, 6 males and 4 females) with no history of neurological or musculoskeletal impairments or disease were enrolled. The study was conducted in accordance with the Declaration of Helsinki and the procedure approved by the Regional Ethics Committee. Informed consent was obtained from all participants after receiving a detailed explanation of the study procedure.

### A. Experimental acquisitions

HDsEMG signals and UUS images were detected simultaneously from the biceps brachii for 8 seconds of low-level (5% of maximum voluntary contraction) isometric elbow flexions (Fig. 1A). The details of the experimental protocol are reported in [7].

A 64-electrode matrix (8 rows by 8 columns with 10 mm inter-electrode distance) designed to allow the simultaneous acquisition of HDsEMG and ultrasound from the same muscle region [14] was placed over the muscle belly. Monopolar EMG signals were detected, conditioned (Bandwidth 10-500 Hz, gain 46 dB), and sampled at 2048 Hz with 16-bit resolution using a wireless HDsEMG acquisition system (MEACS, LISiN, Politecnico di Torino, Turin, Italy) [15].

Radiofrequency (RF) data were acquired using Verasonics Vantage 128 (Verasonics, Inc., Kirkland, WA, USA). Plane-wave data were obtained at a frame rate of 2500 Hz. The L11-5v probe (7.8125 MHz center frequency) was placed over the muscle to obtain cross-sectional scanning between the fourth and fifth rows and was cantered with respect to the columns of the EMG matrix (Fig. 1A).

A rectangular pulse provided by an external generator (StimTrig; LISiN, Politecnico di Torino, Italy) was used to start the ultrasound acquisition and was concurrently acquired by the HDsEMG system to synchronize EMG-ultrasound acquisitions [16].

### B. Data analysis

Experimental signals were processed by implementing both STA and STICA approaches. The two approaches rely on a combined analysis of the firing pattern of active MUs and the tissue velocity sequence estimated from HDsEMG and ultrafast ultrasound imaging, respectively (Fig. 1). Spike trains of individual MU were decomposed from HDsEMG data using a validated method based on convolution kernel compensation (Fig. 1B) [17]. Displacement velocity images were estimated from the RF data using 2D autocorrelation velocity tracking, as in [18], to obtain a sequence of tissue velocities (Fig. 1C). After cropping the images at a depth of ∼ 20 mm (outside the maximum EMG range), the sequence resulted in 19976 frames of 70 × 128 pixels.

#### 1) Spike-triggered averaging (STA) approach

The STA of the tissue velocity sequence was performed using the discharge times of the MU identified from HDsEMG decomposition. For each MU, windows of 125 ms centered on each firing instant were averaged to obtain a short sequence in which the contribution of the considered MU might be emphasized over the others. A single image was extracted from this averaged sequence as follows: (i) the time sample at which the maximum velocity occurred was identified, and (ii) the sequence was averaged across ± 10 frames around the maximum time sample (20 frames in total). A global thresholding at 70% of the maximum was applied to the obtained image, providing the single MU displacement area.

#### 2) Spatiotemporal independent component analysis (STICA) approach

STICA was applied to regions of interest (ROI) of the tissue velocity sequence of 35 × 35 pixels spanning the whole field of view with a shift of 5 pixels in both directions. The details of the method are reported in [7]. Here, a synthetic description is provided. Briefly, the algorithm extracted 50 components for each ROI, optimizing the independence over space and time. The components comprised a temporal signal and a spatial map. For each MU decomposed in HDsEMG, the convolution between its firing discharges and a synthetic waveform, representing the velocity of contracting fibers in the superficial-deep axis, was computed. This convolution produces what we refer to as the train of MU velocity twitches. The cross-correlations between these trains and the temporal signals of all UUS components were computed. For each ROI, the component with the maximum correlation within a ± 20 ms time lag was selected. This procedure provided a set of correlation coefficients for all considered ROIs for each decomposed MU. A group of adjacent ROIs with correlation values > 50% of the maximum of all ROIs was retained. The spatial maps of the ROIs corresponding to the identified group were summed to obtain a single image. A global thresholding at 70% of the maximum was applied to the obtained image, providing the single MU displacement area.

### C. Comparison of STICA and STA approaches

The centroids of the MU displacement areas obtained from STA and STICA were computed and compared with the centroids of the amplitude distribution of the MUAP of the corresponding MU (i.e., the MU whose firings were used as input to STA and STICA algorithms). The comparison was performed by computing the centroid-to-centroid (EMG-ultrasound) distance in the medio-lateral direction.

The two algorithms were processed considering the entire 8 s of acquisition and only the first 2 s of acquisition to evaluate the effect of the signal duration on their performance.

### D. Statistical analysis

The effect of the factors “method” (STA and STICA) and “duration” (2 s and 8 s) were tested with a 2-way ANOVA on the centroid-to-centroid distance. Post-hoc assessments were conducted using the Bonferroni test whenever a main effect was verified. The significance level was set to *p* < 0.05.

## III. Results and Discussion

The HDsEMG decomposition provided 180 MUs (mean ± SD, 18 ± 8 MUs per participant).

Fig. 2 shows the identification of the MU displacement areas of four representative MUs using the combined analysis of HDsEMG data and UUS images. The figure depicts, in the upper panel, the spatial distribution of single differential MUAPs computed longitudinally (i.e., along the fibers’ direction), the regions of displacement in the ultrasound identified with STA (middle panel) and with STICA (bottom panel), respectively. The examples reported in Fig. 2 show four different outcomes of the two algorithms in terms of spatial agreement between the MU displacement area in UUS and the EMG amplitude distribution. Specifically, MU#1 showed a good spatial match for both STA and STICA (as shown by the horizontal distance between the symbols ‘+’ in EMG, ‘?’ for STA and ‘×’ for STICA). MU#2 and MU#3 showed a good match only with STICA and STA, respectively, while the MU displacement area identification seemed to fail for both STA and STICA in MU#4.

**Fig. 2:**
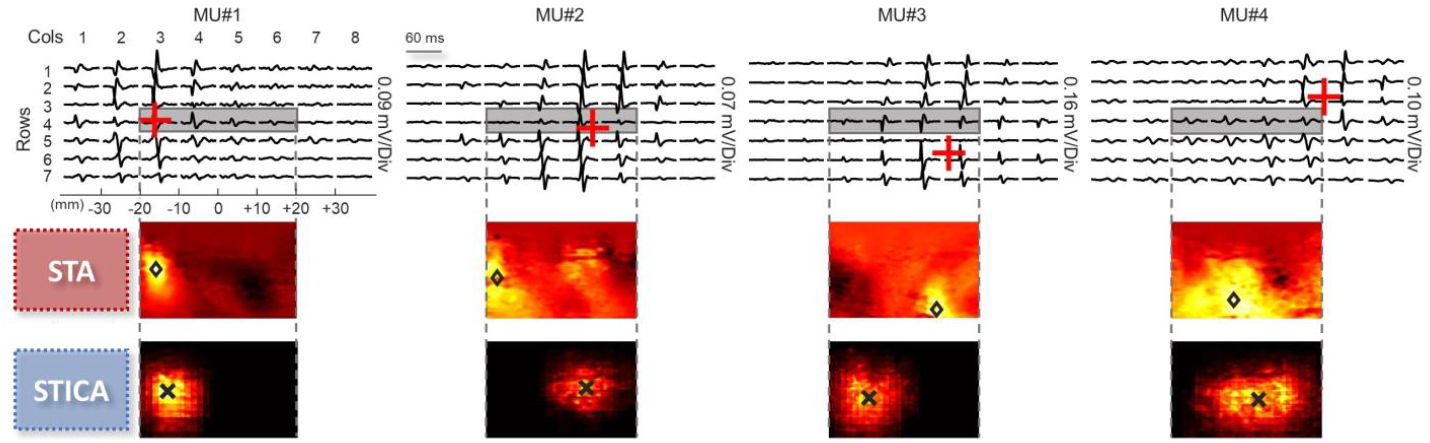
Four examples of MUs (column wise) decomposed from HDsEMG and the corresponding identified region of displacement. First row: template of the motor unit action potential (MUAP) in single differential derivation along the rows of HDsEMG grid. Red + represents the centroid of the MUAP template distribution as calculated in [7]. The grey rectangle shows the region where the ultrasound probe was placed to scan the muscle cross-sectionally. Second row: images obtained with the spike triggered averaging (STA) approach representing the region of displacement of the corresponding MU. ? indicates the centroid of the tissue velocity image. Third row: final images of the approach based on the spatio-temporal independent component analysis (STICA) representing the region of displacement of the corresponding MU. × represents the centroid of the tissue velocity image. The figure shows from left to right: a well-detected MU (low centroid-to-centroid distance between EMG and ultrasound) for both approaches; a MU displacement area correctly identified only by STICA; a MU displacement area correctly identified only by the STA approach; a wrong MU displacement area identification for both STA and STICA.

Fig. 3 shows the boxplots of distances between EMG and ultrasound centroid for all the considered MUs for both STA (red) and STICA (blue) when 8 seconds and 2 seconds of signal duration are considered. STICA approach provided lower distances for all the tested durations, as demonstrated by statistically significant differences with respect to STA (*p* < 0.001). Although not significant (*p* = 0.054), a trend associated with signal duration can be appreciated in both STA and STICA and to a larger extent for STA.

**Fig. 3:**
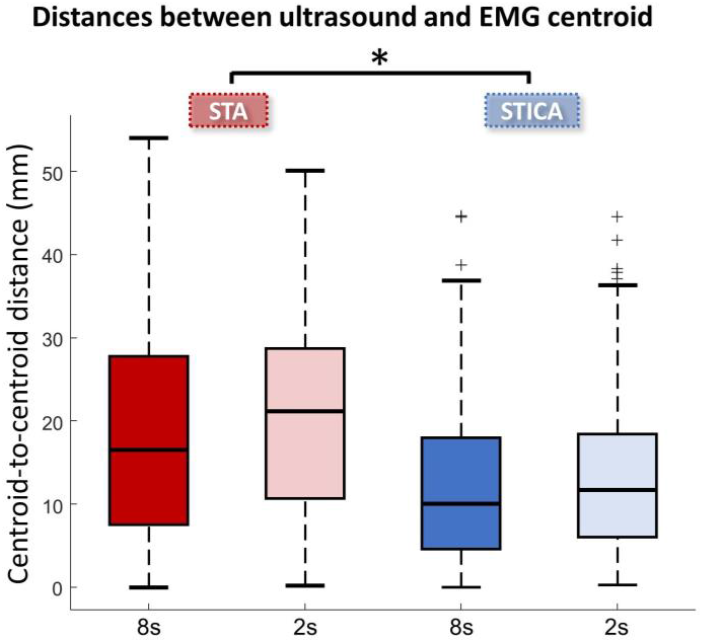
Boxplots of distances between EMG and ultrasound centroid for all the considered MUs for both STA (red) and STICA (blue) when considering 8 seconds (dark color) and 2 seconds (light color) duration of the acquisition. *p<0.001

The distances obtained for STA and STICA were in the range [0–53 mm] and [0–37 mm], respectively and confirmed the results obtained in simulated conditions, with STICA providing lower distances. As expected, these values are larger than those obtained in simulated conditions where distances always lower than 20 mm were obtained by both methods and for contractions up to 20% MVC. Thus, although we considered lower contraction levels (5% MVC), the performance of both methods degraded in experimental conditions considerably. Besides the obvious differences between experimental and simulated conditions (e.g., the effect of connective tissue affecting the transversal force transmission between fascicles), this result can be explained by the fact that the in the experimental analysis, our reference for the “true” medio-lateral location of MU displacement area was the centroid of the MUAP distribution, that can be identified with a relatively low spatial resolution (inter-electrode distance), thus affecting the estimation error.

Concerning the effect of signal length, both approaches were expected to be affected by the duration of the EMG and ultrasound time series. However, our results suggest that STICA may provide better estimations of MU displacement areas when short duration acquisitions are performed (light color boxes in Fig. 3). This can be an important advantage considering the large amount of data associated with the detection of sequences of high frame rate ultrasound images. Moreover, the capability to obtain good displacement area estimations for small signal epochs could allow the implementation of STICA-based algorithms exploiting the temporal segmentation of the detected signal to improve the estimation, e.g., by providing a repeatability index of the identified area across the contraction. On the other hand, the low computational complexity of STA may be preferable for applications where the computational time is relevant.

## IV. Conclusion

This study compared two approaches (STA and STICA) to identify individual MU properties by combining HDsEMG and UUS. We showed that both STICA and STA can be used to detect regions of muscle tissue displacement whose location is associated with the representation of the MU electrophysiological activity (i.e., MUAP distribution). Although wrong estimations occurred for both methods, STICA provides more precise estimation of the MU displacement region location, to the detriment of higher computational complexity. The higher reliability in the assessment of the MU territory provided by STICA may be explained as a result of the decomposition of the sources’ location and temporal information, while STA uses the global interfering signal. Our results expand those of our previous simulation study, providing new qualitative evidence on the suitability of the two proposed approaches for anatomical identification of individual MU using the combination of HDsEMG and UUS analysis. By providing electrophysiological, anatomical and mechanical information on active MUs, these multi-modal approaches are relevant to improve our understanding of the underlying mechanism of muscle contraction, from the neural command to the force production, with applications ranging from basic research to motor rehabilitation.

